# Emergence of alternative stable states in microbial communities in a fluctuating environment

**DOI:** 10.1101/678367

**Authors:** Vilhelm L. Andersen Woltz, Clare I. Abreu, Jonathan Friedman, Jeff Gore

## Abstract

The effect of environmental fluctuations is a major question in ecology. While it is widely accepted that fluctuations and other types of disturbances can increase biodiversity, we have only a limited understanding of the circumstances in which other types of outcomes can occur in a fluctuating environment. Here we explore this question with laboratory microcosms, using cocultures of two bacterial species, *P. putida* and *P. veronii*. At low dilution rates we observe competitive exclusion of *P. veronii*, whereas at high dilution rates we observe competitive exclusion of *P. putida*. When the dilution rate alternates between high and low, we do not observe coexistence between the species, but rather alternative stable states, in which only one species survives and initial species’ fractions determine the identity of the surviving species. The Lotka-Volterra model with a fluctuating mortality rate predicts that this outcome is independent of the timing of the fluctuations, and that the time-averaged mortality would also lead to alternative stable states, a prediction that we confirm experimentally. Other pairs of species can coexist in a fluctuating environment, and again consistent with the model we observe coexistence in the time-averaged dilution rate. We find a similar time-averaging result holds in a three-species community, highlighting that simple linear models can in some cases provide powerful insight into how communities will respond to environmental fluctuations.

## Introduction

In nature, environmental conditions vary over time, and this variation can have significant impacts on the structure and function of ecological communities. Examples of the impacts of environmental variability on community composition include daily cycles of of light and temperature that allow nocturnal and diurnal organisms to coexist, and seasonal variation that causes reproducible succession patterns in communities of plants^1^, freshwater^2^ and marine microbes^3^. Community function can also be strongly influenced by varying environmental conditions. For example, a single rain event can cause up to 10% of annual carbon dioxide production of a forest^4^, due in part to enhanced microbial activity in rewetted dry soil^5^. Varying environmental conditions may even cause ecosystems to abruptly and irreversibly change states, such as lakes that shift from clear to turbid due to human-induced eutrophication^6^ and reefs that transform from kelp forests to seaweed turfs due to heat waves^7^. Given the inevitability of temporal variability in nature, an improved understanding of how this variability affects ecological communities is essential for understanding natural ecosystems.

Both theoretical^8–12^ and empirical^13–17^ studies have shown that disturbances can stabilize communities and enhance diversity. For example, temporal fluctuations of light^18^ and temperature^19^ have been shown to lead to stable coexistence of microbes. One possible mechanism for this effect is that different species are favored at different times, such that no species can ever dominate the system, and the result might be coexistence^20^. For example, species A may exclude species B in one environment, whereas B excludes A in another environment. Fluctuating between the two environments might lead to coexistence of the two species. If true, such an effect could propagate to complex communities with more than two species, leading to more coexistence in the fluctuating environment than in the constant ones. Another possible explanation for coexistence in a fluctuating environment is time-averaging: species A and B might coexist when fluctuating between two environments because they coexist in the constant average environment^21,22^. The former theory predicts that fluctuations are necessary for coexistence, while the latter suggests that coexistence depends upon the average environment, with or without fluctuations.

Compared to a wealth of studies of fluctuation-induced coexistence, less is known about whether perturbations may lead to outcomes other than increased diversity. For example, alternative stable states have been observed in fluctuating environments, such as in gut microbiota communities subject to laxative treatments^23^, intertidal biofilms perturbed by climatic events^24^, and regions of forest and barrens disturbed by frequent fires^25^. In these cases, however, it is often difficult to distinguish between alternative states that are simultaneously stable and different states that are stable in different environments. There is some theoretical support for fluctuation-induced alternative stable states in particular systems^26,27^, but these predictions are challenging to test in the field due to the difficulty of controlling all ecological factors. The proposition that environmental fluctuations may have more complicated effects on ecosystems than simply affecting diversity is therefore in need of systematic study and demonstration.

In this paper, we make use of highly controllable microbial microcosms to explore the effects of temporal fluctuations on communities. We grow bacterial species in liquid culture with daily dilution, and implement environmental fluctuations by alternating the amount of the dilution. The growth-dilution process imposes a tunable death rate on the system, where the dilution factor determines the fraction of cells discarded each day. In a two-species coculture, we observe that fluctuating dilution factors leads to bistability, or two alternative stable states that depend on the initial abundances of each species. To explain this result, we use a simple phenomenological model: the Lotka-Volterra competition model adapted to incorporate a fluctuating global mortality rate. This model with a fluctuating mortality rate predicts that the bistability is independent of the timing of the fluctuations, and that the time-averaged mortality would also lead to alternative stable states, a prediction that we confirm experimentally. The model predicts that fluctuating mortality can result not only in bistability, but also in stable coexistence, depending on the strength of interspecies inhibition, which we confirm experimentally. More broadly, the model predicts that an environment with a fluctuating death rate equilibrates to the same outcome as the time-averaged added mortality rate. We test this prediction both in two-species cocultures and in a more complex community with three species, where a fluctuating death rate and a constant death rate lead to the same qualitative outcome. These results suggest that fluctuations can in some cases have predictable consequences on community structure.

## Results

To explore the effect of a fluctuating environment on an experimentally tractable microbial community, we performed coculture experiments with fluctuating as well as constant dilution factors. The cultures were allowed to grow for 24 hours and then diluted by transferring a small amount of culture into fresh growth media. Because the total experimental volume remains constant, the amount of culture added from the previous day determines the dilution factor. We began by coculturing *Pseudomonas veronii* (*Pv*) and *Pseudomonas putida* (*Pp*). At a low dilution factor (10×), *Pv* competitively excludes *Pp*, as the fraction of *Pp* goes to zero from all initial fractions (Fig. 1A). At a high dilution factor (10^4^×), the outcome is reversed, with *Pp* excluding *Pv* as its fraction goes to one from all initial fractions (Fig. 1B). We refer to both of these outcomes as competitive exclusion because the final state does not depend on starting conditions; all starting fractions move toward one stable state, which is either zero or one.

**Figure 1:**
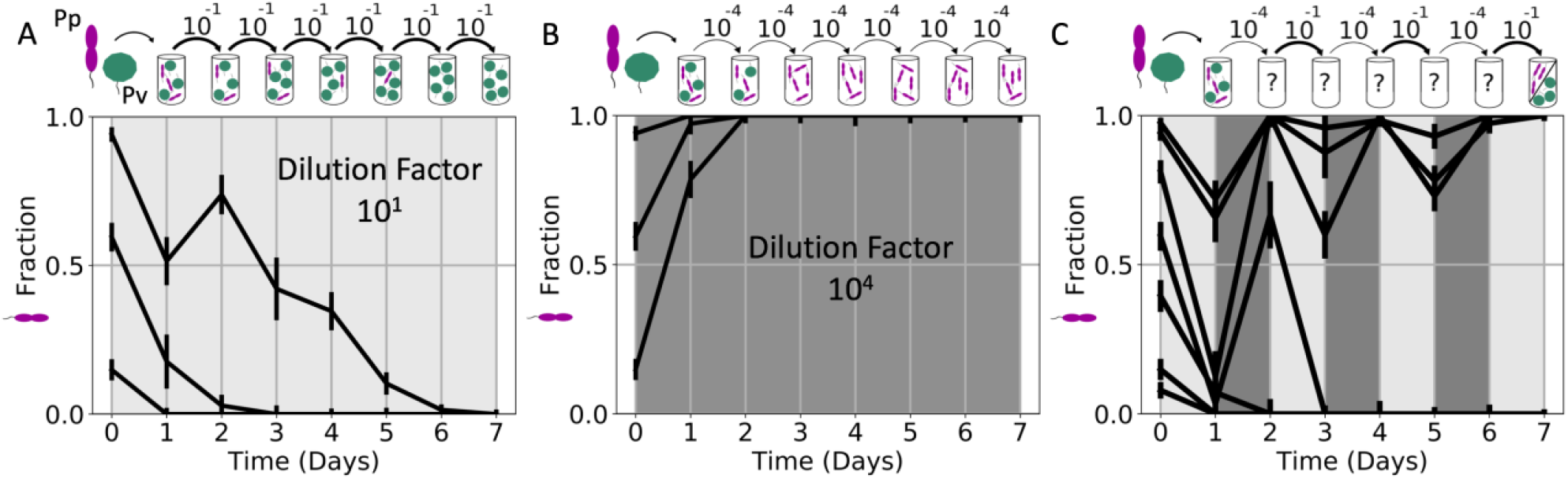
Experimental observation of alternative stable states in a fluctuating environment. **A**: When a coculture of Pp (purple) and Pv (green) was diluted by a factor of 10 each day (1/10 of the previous day’s culture transferred to fresh media, keeping the volume constant), the slow-growing Pv dominated, sending the fraction of fast grower Pp to zero from several starting fractions. **B**: When the same culture was subject to a much higher dilution factor, 10^4^, fast grower Pp dominated. **C**: Surprisingly, fluctuating between the low and high dilution factors shown in A and B resulted in alternative stable states. Either Pp or Pv can dominate, depending on their relative initial abundances. In all plots, we qualitatively indicate the dilution factor for that day by the shading of the plot; low dilution factors have a lighter shading, while high dilution factors have a darker shading. Error bars are the SD of the beta distribution with Bayes’ prior probability (see Methods).

Given that there was a single stable state at low dilution and a single stable state at high dilution, we expected that alternating between the two dilution factors would also lead to a single stable state (which could be survival of just one species or possibly stable coexistence of the two). To our surprise, when we performed the coculture experiment alternating between the two dilution factors, we instead observed the emergence of alternative stable states—only a single species survived, but the surviving species depended on the species’ initial fractions (Fig. 1C). We therefore observe bistability in the fluctuating environment despite the fact that neither environment alone showed bistability.

To explain a possible origin of this emergent bistability, we employed the Lotka-Volterra (LV) competition model with added mortality, which previously provided powerful insight into how microbial competitive outcomes shift with dilution rate^28^. The two-species LV model with added mortality is:

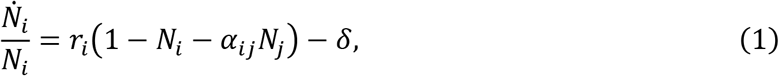

where *N*_*i*_ is the abundance of species *i* normalized by its carrying capacity, *r*_*i*_ is the maximum growth rate for species *i*, *α*_*ij*_ is the competition coefficient that determines how strongly species *j* inhibits species *i*, and *δ* is the imposed mortality rate, which is experimentally controlled by the dilution factor, which specifies the fraction of cells discarded per day. In the absence of added mortality *δ*, the outcome is independent of growth rates *r*_*i*_ and solely determined by whether the competition coefficients *α*_*ij*_ are greater or less than one: coexistence and bistability result when both are less than or greater than one, respectively, and dominance results when only one coefficient is greater than one (Fig. 2B). The presence of mortality makes the competition coefficients, and thus the outcome, functions of growth and mortality, as can be seen in the reparameterized model (Supplementary Note 1):

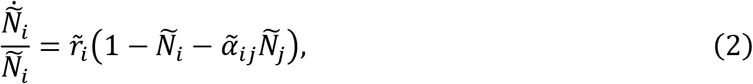

where

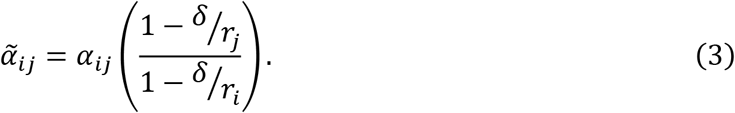

**Figure 2:**
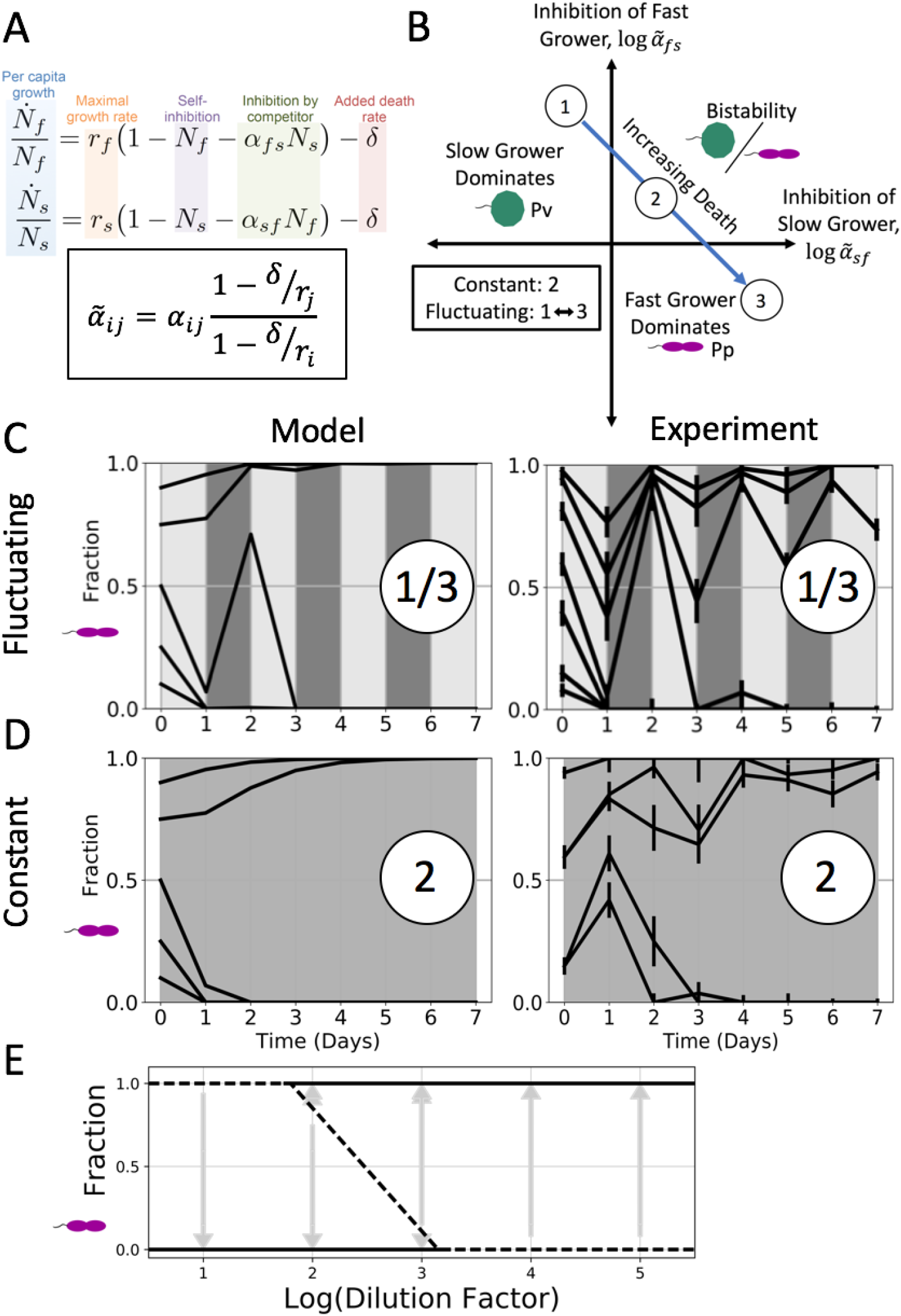
Bistability occurs in both fluctuating and average environments, confirming model prediction. To understand the results from Fig. 1, we employed a modified Lotka-Volterra model. **A**: We model daily dilutions by adding a community-wide death rate term *δ* to the two-species Lotka-Volterra (LV) competition model. The per-capita growth rate is a function of a species’ maximum growth rate, self-inhibition, and competition with the other species. Because the per capita growth rate is linear and additive, the model predicts that the outcomes of a fluctuating environment should be the same as that of the time-averaged environment (Supplementary Note 1). **B**: The solutions to the LV model can be represented by a phase space of the re-parameterized competition coefficients ⍺, which are functions of death *δ* and growth *r*. If a slow grower dominates at low or no added death, increasing mortality will favor the fast grower, causing the pair to pass through a region of bistability or coexistence on the way to fast grower dominance. Here we have illustrated the trajectory of a bistable pair. **C-D**: To test the prediction that the fluctuating and time-averaged environments are qualitatively equivalent, we cocultured Pp and Pv at a dilution factor equal to the time-average of the fluctuating dilution factors in Fig. 1, and we observed bistability, confirming this prediction (lower right). Additionally, we simulated daily dilutions of the model with both constant (upper left) and fluctuating (lower left) dilution factors and observed good agreement between the model and experimental results. **E**: A bifurcation diagram of the Pp-Pv outcomes at all constant dilution factors shows that the fast-growing Pp is favored as dilution increases. Arrows represent time trajectories from initial to final fractions; solid lines represent stable equilibria, and dotted lines represent unstable fractions. This diagram was used to estimate the competition coefficients used in simulations (see Methods and Supp. Fig. 1). Error bars are the SD of the beta distribution with Bayes’ prior probability (see Methods).

If a slow grower dominates at low mortality/dilution, the model predicts that increasing dilution will reverse the outcome and cause the fast grower to win, and at some range of intermediate dilution the pair passes through either a region of bistability or coexistence (Fig. 2B). Moreover, a fluctuating dilution rate will lead to the same outcome as the time-averaged rate in the absence of fluctuations. This prediction arises because the per-capita growth rates, 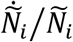, in the LV model are linear and additive, and the steady state reached through a temporally fluctuating mortality *δ* is the same state reached by its linear time-average ⟨*δ*⟩ (Supplementary Note 2). The LV model thus makes the simple prediction that an experiment alternating between dilution factors 10 and 10^5^, for example, will end in the same state as one with a constant dilution factor of 10^3^. When averaging dilution factors, we use the geometric mean of the dilution factors because of the logarithmic relation between discrete dilution factor and equivalent continuous rate *δ* (Supplementary Note 2).

Given our experimental observation of bistability in a fluctuating environment (Fig. 1), our next step was to look for bistability in the constant average environment, as the model predicts. We cocultured *Pp* and *Pv* at a range of constant and fluctuating daily dilution factors. The fluctuating dilution factor experiments again led to bistability, replicating our previous results (Fig. 2C-D). The constant dilution factor experiments revealed a range of fixed points varying with dilution factor: in addition to domination of slow-growing *Pv* at low dilution and that of fast-growing *Pp* at high dilution, we observed bistability at two intermediate dilution factors (Fig. 2E). The separatrix, or starting fraction dividing the two stable states, shifts in favor of the fast grower at the higher of these two dilution factors, consistent with the model. These experimental results confirm the model’s prediction that the dilution factor can be time-averaged to result in the same qualitative outcome; namely, bistability (see Supp. Fig. 3).

As previously mentioned, the LV model also predicts that pairs of species can coexist at intermediate mortality rates. Given this additional prediction, we sought to experimentally verify that coexistence can also result from time-averaging the dilution factors. Based on results of previous cocultured experiments in a similar growth medium^28^, we chose another fast grower, *Enterobacter aerogenes* (*Ea*), which is also excluded by the slow-growing *Pv* at low dilution factor. At intermediate dilution factor *Ea* and *Pv* coexist, and *Ea* dominates at high dilution factor (Fig. 3D). We then fluctuated between the dilution factors in which either species dominated, as in our initial experiments. This time, we observed stable coexistence in the fluctuating environment (Fig. 3). Furthermore, the stable fraction of *Ea* fluctuated in the neighborhood of the stable fraction of Ea in the constant dilution experiment. In both the bistable pair (Fig. 2C-D) and the coexisting pair (Fig. 3B-C), simulations match the experimental trajectories over time, in both constant and fluctuating dilution factors. These examples of agreement between the model and experiments emphasize that in our simple bacterial communities, a fluctuating dilution factor may be time-averaged (see Supp. Fig. 4).

**Figure 3:**
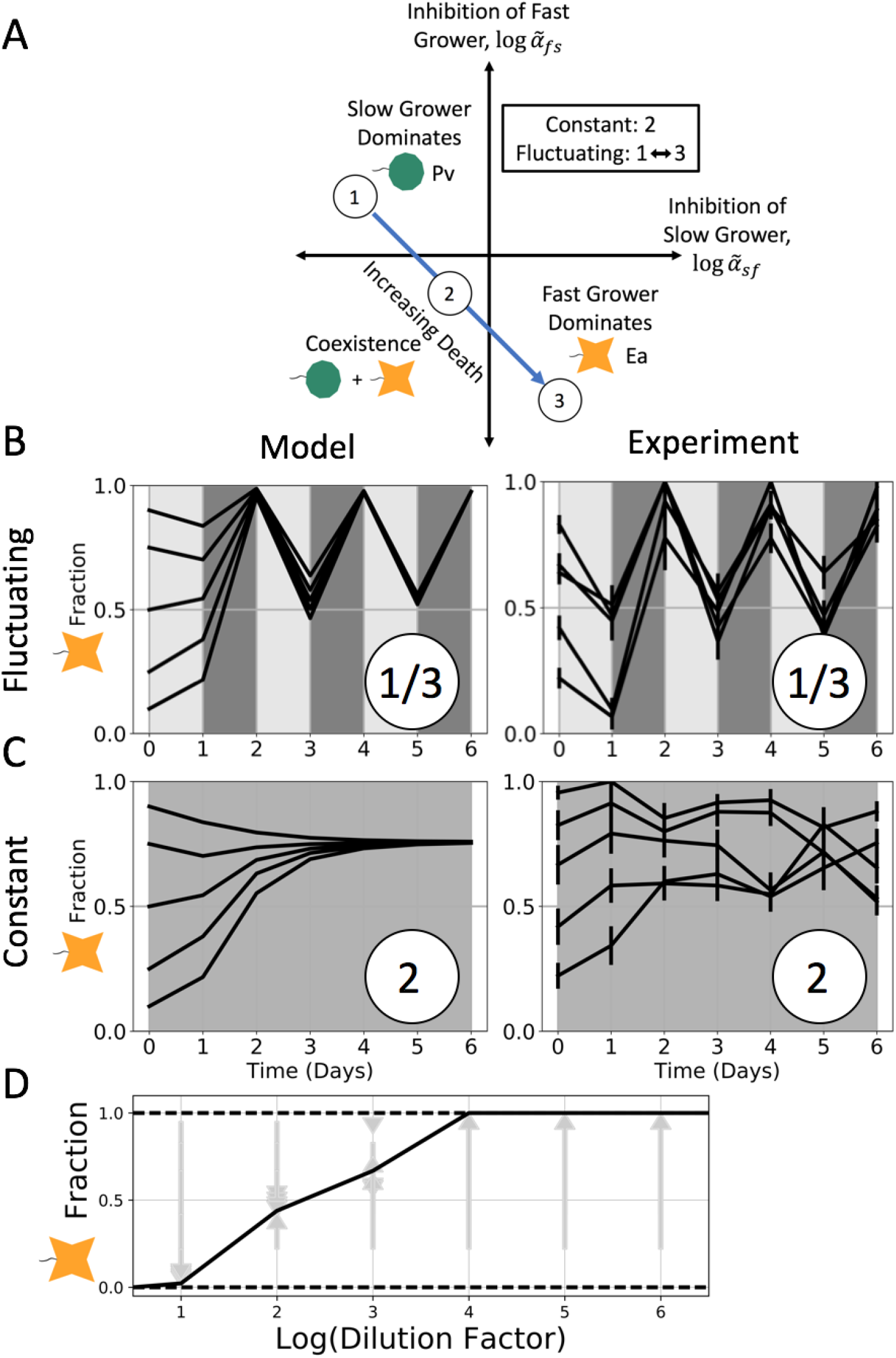
Coexistence occurs in both fluctuating and average environments, confirming model prediction. **A**: As in Fig. 2, the LV model phase space shows qualitative outcomes divided by the zero-points of the logarithm of the re-parameterized competition coefficients. Here we have illustrated the trajectory of a pair that passes through the coexistence region as the death rate increases. **B-C**: When we cocultured slow-growing Pv with another fast grower, Ea, we observed coexistence in an environment that fluctuated between dilution factors in which either species dominated (upper right). At a dilution factor equal to the mean of fluctuations, we also observed coexistence (lower right), confirming the model’s prediction about a time-averaged environment for both types of trajectories across the phase space. Once again, simulations of daily dilutions showed good agreement between the model and experimental results (left). **D**: A diagram of outcomes at all constant dilution factors shows that fast-growing Ea is favored as dilution increases. Arrows represent time trajectories from initial to final fractions; solid lines represent stable equilibria, and dotted lines represent unstable fractions. We used this diagram to estimate the competition coefficients used in simulations (see Methods and Supp. Fig. 2). Error bars are the SD of the beta distribution with Bayes’ prior probability (see Methods).

While pairwise interactions in a fluctuating environment appear to be well-described by the LV model, the same might not be true of more complex communities. To address this question, we cocultured all three of the previously mentioned species, *Ea*, *Pp*, and *Pv*. Across a range of constant dilution factors, we saw changes in community outcome as a function of the dilution factor (Supp. Fig. 6), shifting from a *Pv*-dominated final state at low dilution to coexistence of *Pp* and *Ea* at high dilution. At an intermediate dilution factor, bistability occurs in the trio between a *Pp*-*Ea* coexistence state and a *Pv*-*Ea* coexistence state (Fig. 4A). This bistability of coexisting states can be predicted from the corresponding pairwise results, using our previously developed community assembly rules^29^. When we fluctuated the dilution factor between the low and high values that average to this intermediate dilution factor, we again observed bistability, with each initial condition ending in the same final state as it did in the constant environment (Fig. 4B). These results show that even in a more complex community of three species, a fluctuating mortality rate still leads to the same results as the constant average environment, which is in line with the predictions of the modified LV model.

**Figure 4:**
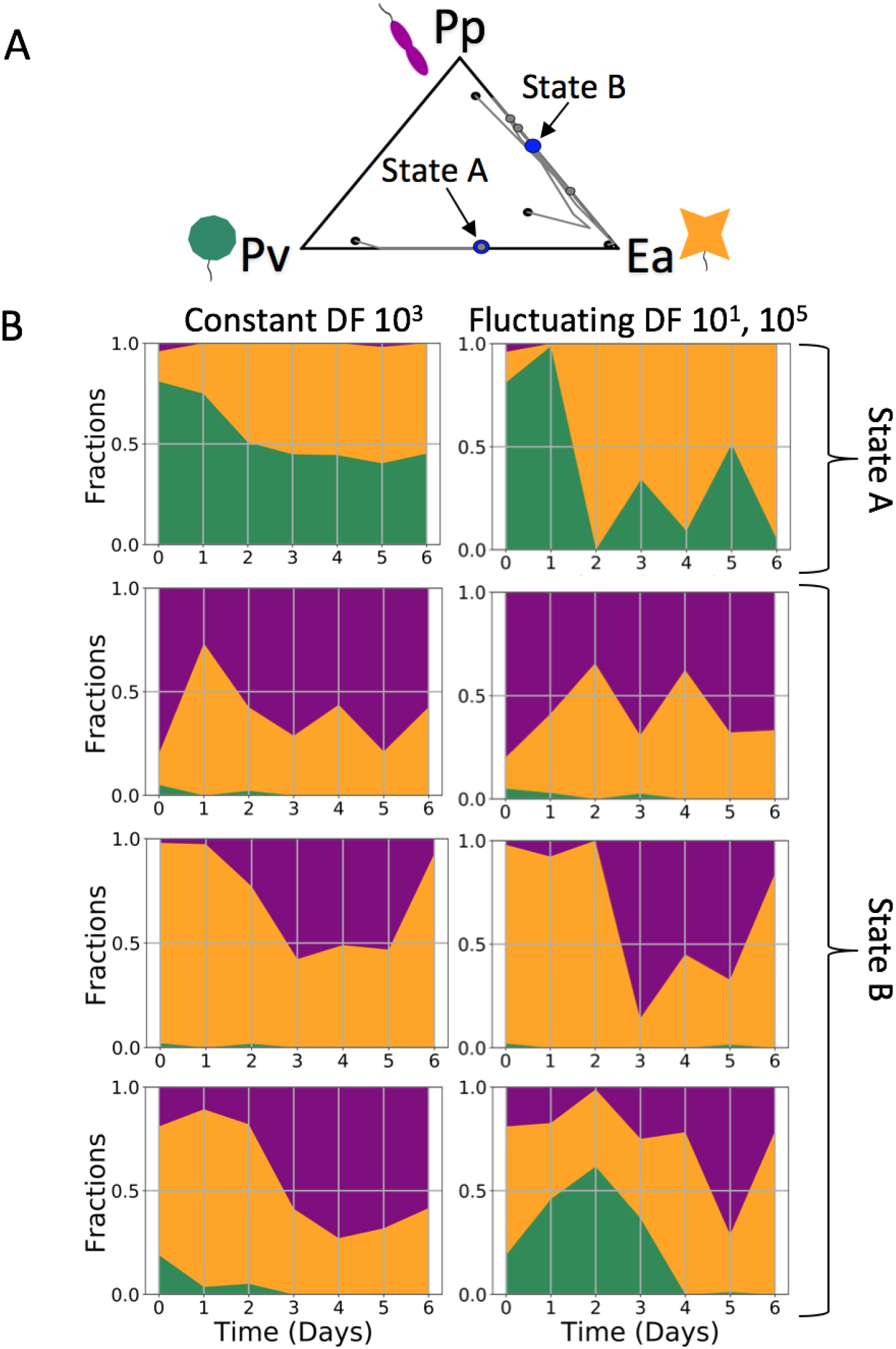
Fluctuating environment predictably leads to alternative stable states in a three-species community. To ensure that the time-averaging prediction is not only applicable to simple two-species communities, we tested our results in a three-species environment. **A**: A ternary plot shows the outcomes of a coculture with all three species, Pv, Pp, and Ea, in a constant environment with dilution factor (DF) 10^3^. The time trajectories, indicated by the grey lines, end at one of two alternative stable states (shown in blue; for state B, we plot the average of the three trajectories). In state A, Pv and Ea coexist, while Pp and Ea coexist in state B. **B**: Time series plots show the results of three-species coculture experiments in both constant (DF 10^3^; left column) and fluctuating (between DF 10^3^ and DF 10^5^; right column) environments. As indicated in **A**, three initial fractions end in state B and one ends in state A. We find that the outcomes of a given initial fraction go to the same final state in both the constant and fluctuating environments. This suggests that the ability to time-average the outcomes extends to communities with more than two species (see Supp. Fig. 6).

## Discussion

In criticizing the use of generic hypotheses, JW Fox posited in 2013, “The time is ripe for ecologists to … embark on a new research program testing the assumptions and predictions of logically valid models of diversity and coexistence in fluctuating environments.”^21^ Our results highlight a clear experimental demonstration of a simple theoretical prediction about the effects of fluctuations on community structure. Using simple microbial communities with up to three species, we have shown that an experiment with a dilution factor fluctuating between two extremes, both of which typically lead to competitive exclusion, in fact leads to either coexistence or bistability. The outcome of the fluctuating dilution factors is the same as that of the equivalent time-averaged dilution factor, and therefore appears to be independent of fluctuations. Whether the community coexists or forms alternative stable states in the fluctuating environment can be predicted by the community state in a constant environment, providing evidence that a fluctuating environment can be similar to the constant mean environment.

That the outcome of a fluctuating environment would be the same as that of the equivalent constant environment is consistent with models that have linear and additive per-capita growth rates^21,30^. The Lotka-Volterra competition model, modified to include an added mortality rate, is one such model, which we used for guidance. Other ecological competition models, such as those that explicitly define resource consumption with the Monod equation, are not linear or additive, and do not make this prediction^31^. Although our results rely on what some may call simplistic predictions, our conclusions are strengthened by the simplicity of both the experimental system and the model. More complex empirical studies that make no predictions about the effect of fluctuations, or which seek to (dis)prove a generic prediction such as the intermediate disturbance hypothesis (IDH)^13,14,32–34^, lack clearly defined predictions and thus make less clear conclusions. The IDH, which predicts a diversity peak at some undefined range of intermediate disturbance frequencies, has left some unconvinced^21^, and a meta-study of the phenomenon found little evidence of the trend^35^. This paper provides a case study for how the effect of fluctuations can be understood without adhering to any particular hypothesis of disturbance-diversity relationships, with guidance from a simple phenomenological model.

The absence of resources from the LV model is not its only simplification—it also assumes perfect logistic growth and interactions that are not density-dependent. While these assumptions are clearly simplifications of the dynamics present within any actual community, the model nonetheless worked well in predicting the outcome of fluctuations in the dilution rate in our microbial community. We used regularly alternating and moderate disturbances, but as the duration and intensity of disturbances increases one expects that eventually the model’s predictions will fail due to stochastic extinction caused by finite population sizes. Still, its success here highlights the relevance of simple, phenomenological models to biological systems.

Environmental disturbances have long been thought to weaken competition, thus leading to increased biodiversity^36^. This logic is oversimplified^22^, however, and has inhibited exploration of other potential outcomes of fluctuations, such as alternative stable states. In our experiments, we interpreted alternative stable states to be the result of simple time-averaging of dilution factors, meaning that the outcome, whether bistability or coexistence, did not depend on fluctuations per se. In the LV model, dilution *δ* can be time-averaged because it does not covary with other quantities. Not all types of fluctuations would meet this criteria, however; for example, a disturbance affecting the LV competition coefficients *α*_*ij*_ would lead to a covariance between the coefficients and species’ abundances. As noted previously, temperature fluctuations led to stable coexistence of two species of microalgae^19^, and in that case the outcome differed from the outcome at the mean temperature. Indeed, a version of the LV model that assumes temperature acts only upon growth rates *r*_*i*_ does not allow for linear time-averaging (Lax et. al., pre-print). A question for future investigation is whether there are other simple examples of fluctuating parameters that can be time-averaged.

The effect of environmental disturbances on community structure has become more urgent as many habitats face the effects of climate change. The question of which types of disturbances can be time-averaged is one that may provide insight into the effects of environmental fluctuations on natural communities. Our system was composed of three species of soil bacteria, raising the question of whether communities with more than three species would be similarly affected. A linear time-dependence of interactions and growth rates on added mortality may be more dominant in simple communities, while higher-order effects may become more important in complex communities. Additionally, our simple community results do not mean that there were no nonlinearities or covariances in our system, but only that they were not sufficiently large so as to alter the experimental outcome. Many studies of natural systems have found evidence of nonlinearities and covariances, such as the storage effect^37–39^. In such systems, it may be difficult to disentangle which effects are independent of fluctuations, but our results argue that in some cases, simple models retain their predictive power even in fluctuating environments.

## Methods

### Species and media

The soil bacterial species used in this study were *Enterobacter aerogenes* (Ea, ATCC#13048), *Pseudomonas putida* (Pp, ATCC#12633) and *Pseudomonas veronii* (Pv, ATCC#700474). All species were obtained from ATCC. All coculture experiments were done in S medium, supplemented with glucose and ammonium chloride. It contains 100 mM sodium chloride, 5.7 mM dipotassium phosphate, 44.1 mM monopotassium phosphate, 5 mg/L cholesterol, 10 mM potassium citrate pH 6 (1 mM citric acid monohydrate, 10 mM tri-potassium citrate monohydrate), 3 mM calcium chloride, 3 mM magnesium sulfate, and trace metals solution (0.05 mM disodium EDTA, 0.02 mM iron sulfate heptahydrate, 0.01 mM manganese chloride tetrahydrate, 0.01 mM zinc sulfate heptahydrate, 0.01 mM copper sulfate pentahydrate), 0.93 mM ammonium chloride, 1 mM glucose. 1X LB broth was used for initial inoculation of colonies. Plating was done on rectangular Petri dishes containing 45 ml of nutrient agar (nutrient broth (0.3% yeast extract, 0.5% peptone) with 1.5% agar added), onto which diluted 96-well plates were pipetted at 10 ul per well.

### Growth rate measurements

Growth curves were captured by measuring the optical density of monocultures (OD 600 nm) in 15-minute intervals over a period of ~40 hours (Fig. S3). Before these measurements, species were grown in 1X LB broth overnight, and then transferred to the experimental medium for 24 hours. The OD of all species was then equalized. The resulting cultures were diluted into fresh medium at factors of 10^−8^ to 10^−3^ of the equalized OD. Growth rates were measured by assuming exponential growth to a threshold of OD 0.1, and averaging across many starting densities and replicates (n = 16 for all species).

### Coculture experiments

Frozen stocks of individual species were streaked out on nutrient agar Petri dishes, grown at room temperature for 48 h and then stored at 4 °C for up to two weeks. Before competition experiments, single colonies were picked and each species was grown separately in 50 ml Falcon tubes, first in 5 ml LB broth for 24 h and next in 5 ml of the experimental media for 24 h. During the competition experiments, cultures were grown in 500 μl 96-well plates (BD Biosciences), with each well containing a 200-μl culture. Plates were incubated at 25°C and shaken at 400 rpm, and were covered with an AeraSeal film (Sigma-Aldrich). For each growth–dilution cycle, the cultures were incubated for 24 h and then serially diluted into fresh growth media. Initial cultures were prepared by equalizing OD to the lowest density measured among competing species, mixing by volume to the desired species composition, and then diluting mixtures by the factor to which they would be diluted daily (except for dilution factor 10^−6^, which began at 10^−5^ on Day 0, to avoid causing stochastic extinction of any species). Relative abundances were measured by plating on nutrient agar plates. Each culture was diluted in phosphate-buffered saline prior to plating. Multiple replicates were used to ensure that enough colonies could be counted. Colonies were counted after 48 h incubation at room temperature. The mean number of colonies counted, per plating, per experimental condition, was 49. During competition experiments, we also plated monocultures to determine whether each species could survive each dilution factor in the absences of other species. *Pv* went extinct in the highest two dilution factors (10^−5^ and 10^−6^); other species survived all dilution factors.

### Estimating competition coefficients for simulations

In order to simulate the effect of a fluctuating environment on the pairs (Fig. 2C-D, Fig. 3B-C), we measured growth rates and carrying capacities (Supp. Fig. 5) and estimated the competition coefficients *α*_*ij*_. We used the diagrams of pairwise coculture outcomes at all constant dilution factors (Fig. 2E, Fig. 3D) to estimate the competition coefficients as follows. The different outcomes (dominance/exclusion, coexistence, and bistability) are divided by the zero points of 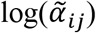 and 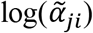, meaning that the qualitative pairwise outcome changes at a dilution factor where one of the reparamaterized coefficients 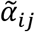 is equal to one. We used Eqn. 3 of the main text to solve for the competition coefficient at the boundary dilution factor where unstable (dotted) and stable (solid) lines intersect on the diagrams (Fig. 2E, Fig. 3D).

### Statistical analysis

The p-values given in Supplementary Figures 5 and 6 were obtained using two-tailed t-tests. The error bars shown in the time-series plots in Fig. 1, Fig. 2, and Fig. 3 are the SD of the beta distribution with Bayes’ prior probability:

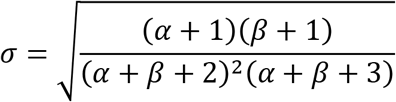

Here, *α* and *β* are the number of colonies of two different species.

### Code availability

Code for data analysis is available upon request.

### Data availability

The source data underlying all figures are provided as a Source Data File. Access to the data is also publicly available at TBD. A reporting summary for this Article is available as a Supplementary Information file.

