## Supplementary Figures and Information for "Emergence of alternative stable states in microbial communities in a fluctuating environment"

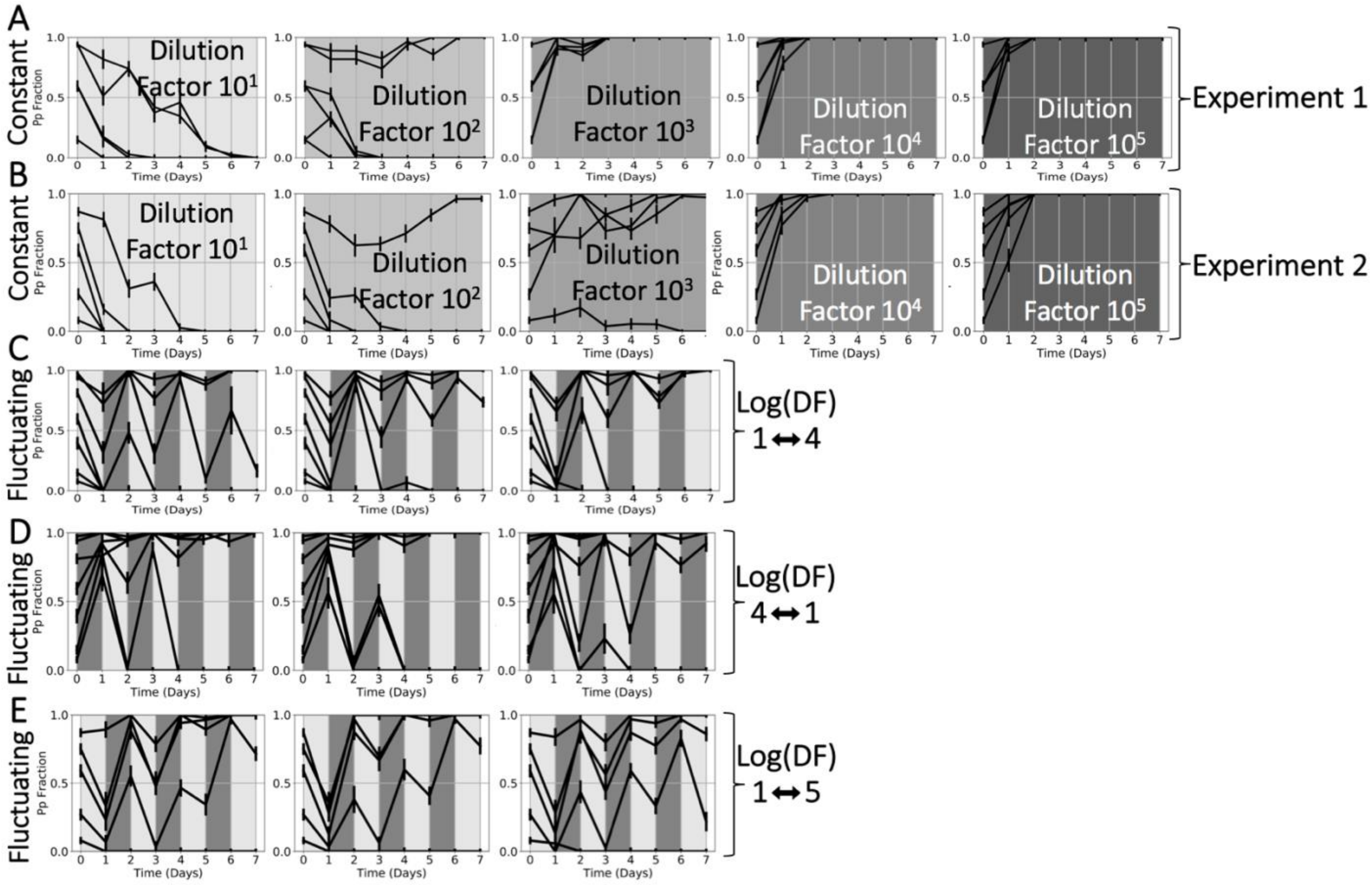


**Supplementary Figure 1: Bistability in pair Pp-Pv is reproducible in both constant and fluctuating environmental conditions.** **A-B**: The data used to generate Fig. 2E of the main text shows that Pp-Pv is a reproducibly bistable pair at dilution factor (DF) $10^2$, and is sometimes bistable at DF $10^3$, although the lowest starting fraction in Experiment 2 might have been above the separatrix. In both cases, the separatrix changes slightly between experiments. Slow-growing Pv is reproducibly dominant at DF $10^1$, and fast grower Pp is dominant above DF $10^3$. Two technical replicates of three starting fractions are shown in **A**, and a second biological replicate of five starting fractions is shown in **B**. **C-E**: Alternative stables states form in a fluctuating environment, and the outcome trends toward that of the average DF ($10^{2.5}$ in panels **C-D**, $10^3$ in panel **E**). Note that the starting fractions near the separatrix (~0.6-0.8 in panels **C-D**, ~0.1-0.3 in panel **E**) take the longest to equilibrate, and may need longer than seven days to reach an absorbing boundary. Each plot shows one technical replicate of seven starting fractions. Panels **C** and **D** show data sampled from the same biological replicate as panel **A**; panel **E** from the same biological replicate as panel **B**. Error bars are the SD of the beta distribution with Bayes' prior probability (see Methods).

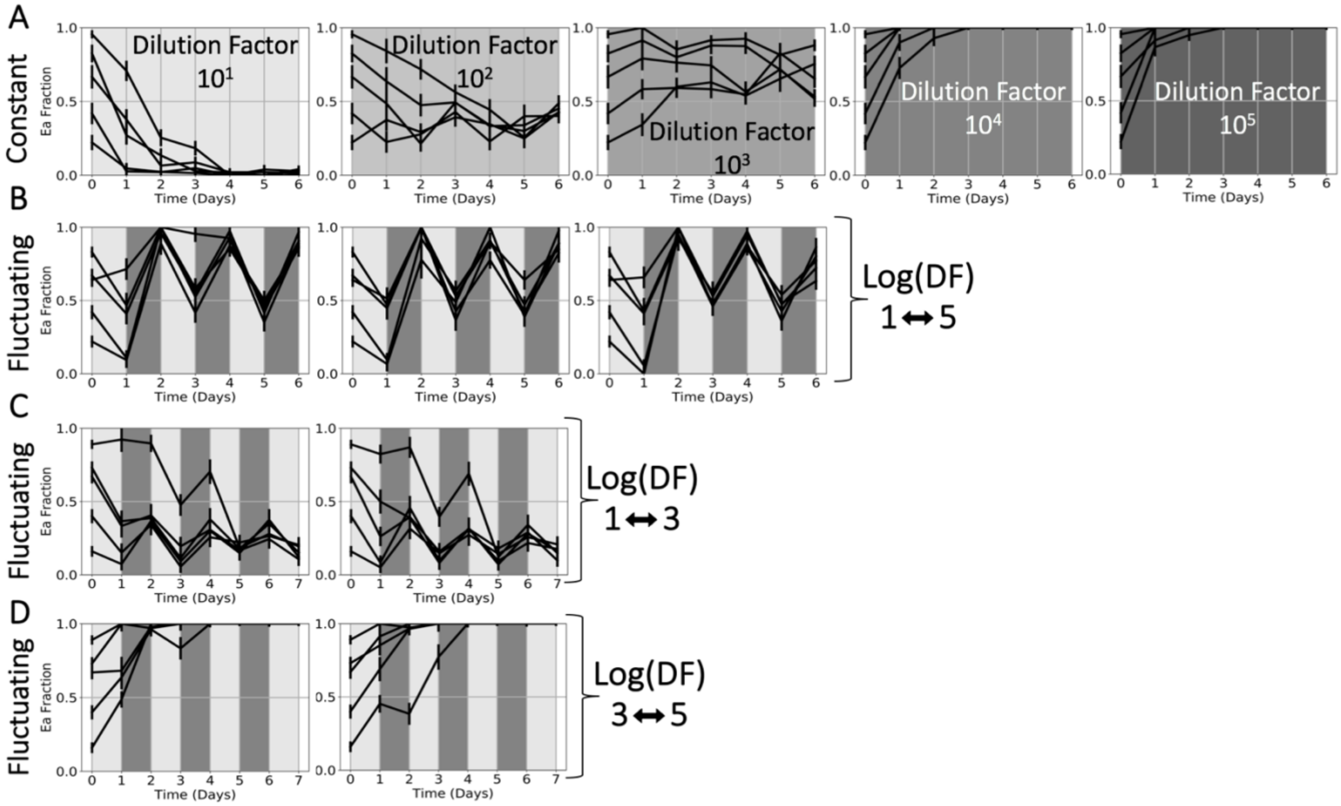


**Supplementary Figure 2: Coexistence in pair Ea-Pv is reproducible in both constant and fluctuating environmental conditions.** **A**: The data used to generate Fig. 3D of the main text shows a reproducible shift as dilution factor (DF) increases, from dominance of slow grower Pv to coexistence to dominance of fast grower Ea. Each plot shows one technical replicate of five initial fractions. **B-D**: Coexistence results in a fluctuating environment if it also results in a constant environment subject to the average DF ($10^3$ in panel **B**, $10^2$ in panel **C**). Furthermore, the coexisting fractions in the fluctuating environments match those of the constant environments. On the other hand, competitive exclusion of Pv results in a fluctuating environment when it also results in the constant DF ($10^4$, panel **E**). Each plot shows one technical replicate of five initial fractions. Panels **C** and **D** were sampled from a different biological replicate than panels **A** and **B**. Error bars are the SD of the beta distribution with Bayes' prior probability (see Methods).

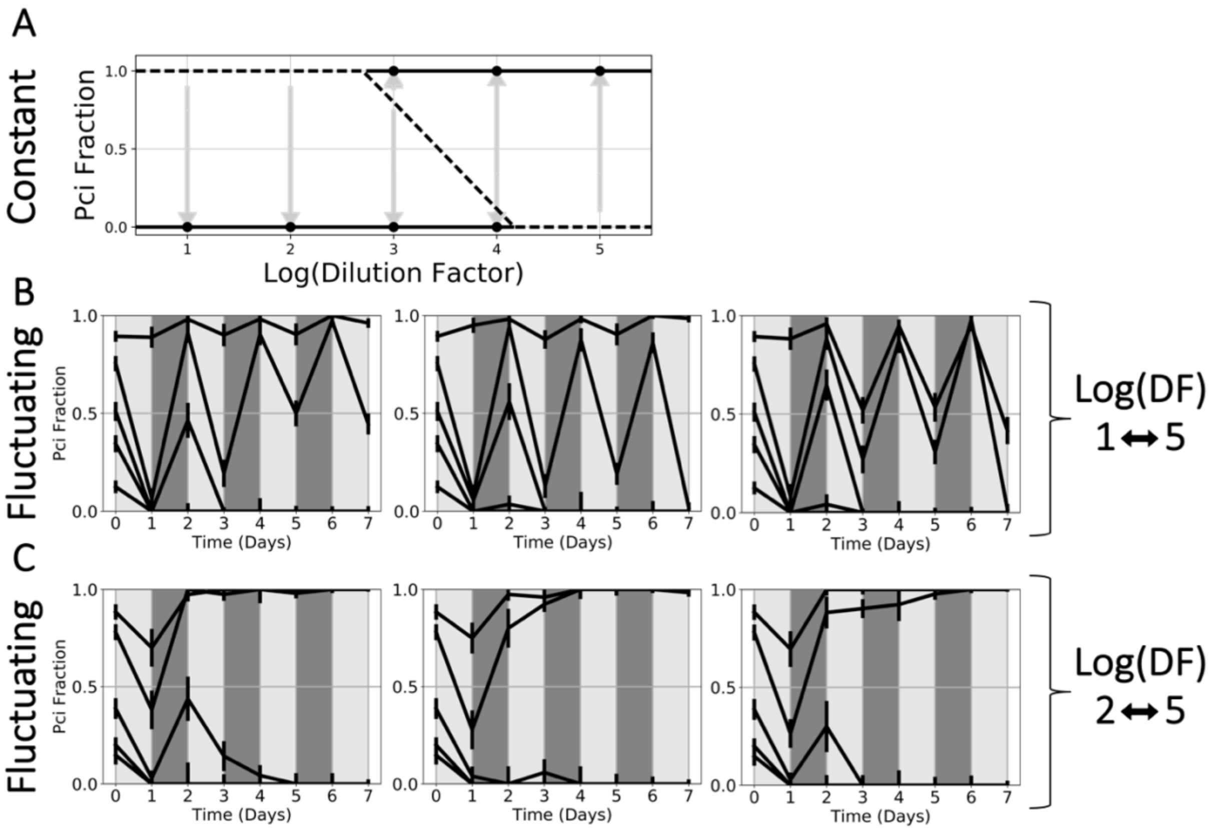


**Supplementary Figure 3: Bistability in additional pair (*Pseudomonas citronellolis* (Pci)-Pv) is reproducible in both constant and fluctuating environmental conditions..** We conducted experiments with another pair found to exhibit alternative stable states, as can be seen from the bifurcation diagram in **A**, which includes data from two different biological replicates. **B**: In an environment fluctuating between DF $10^1$ and DF $10^5$, most trajectories reach the absorbing boundary of zero; the highest initial fraction of Pci is very close to the separatrix in the equivalent constant environment (DF $10^3$) and as such does not consistently go to a single final outcome. One should note that these results do not violate the time-averaging prediction of the LV model, since perturbations near a separatrix may cause a trajectory to cross the separatrix and thus take longer to reach equilibrium. Each plot shows one technical replicate of five initial fractions. **C**: The outcome is more predictable when we fluctuate between DF $10^2$ and $10^5$ for a constant equivalent environment of DF $10^{3.5}$, in which the estimated separatrix is about midway between the absorbing boundaries of one and zero. Here we see alternative stable states forming in the fluctuating environment more clearly, depending on whether a starting fraction is closer to one or zero. Each plot shows one technical replicate of five initial fractions, all of which were drawn from a different biological replicate than in panel **B**. Error bars are the SD of the beta distribution with Bayes' prior probability (see Methods).

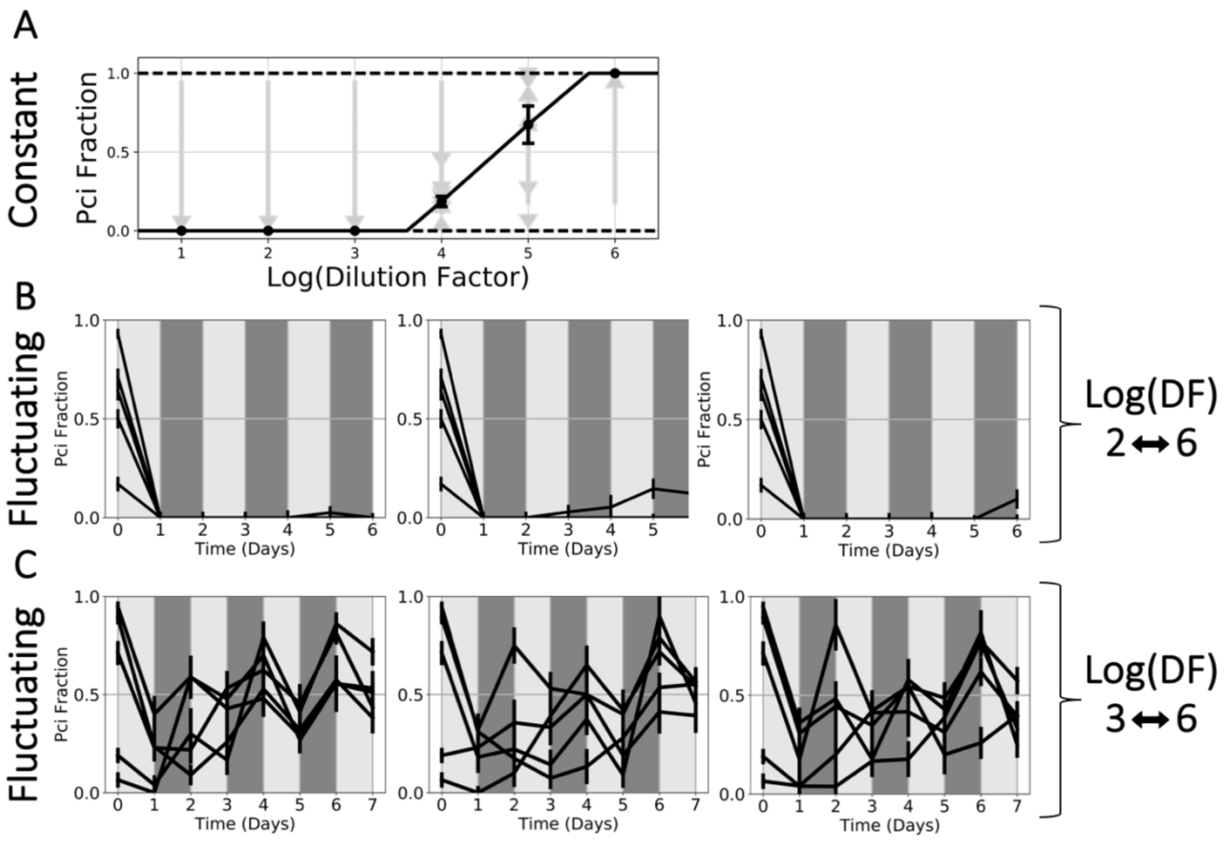


**Supplementary Figure 4: Coexistence in additional pair (Pci-*Pseudomonas aurantiaca* (Pa)) is reproducible in both constant and fluctuating environmental conditions.** We conducted experiments with another pair found to coexist, as can be seen from the bifurcation diagram in **A**, which includes data from two different biological replicates. Error bars are the SEM of all replicates (n=6; 2 biological replicates of 3 starting fractions each). **B:** In an environment fluctuating between DF $10^2$ and DF $10^6$, for a constant equivalent environment at DF $10^4$, the model predicts coexistence at a stable fraction of Pci of ~0.2, as seen in **A**. The failure of coexistence here may be due to stochastic extinction, or domination by Pa in the first 24-hour cycle. Neither of these occurrences would violate the time-averaging prediction of the model. Each plot shows one technical replicate of five initial fractions. **C**: In an environment fluctuating between DF $10^3$ and $10^6$ for a constant equivalent environment of $10^{4.5}$, we expect a stable coexisting fraction further from exclusion. As such, we more clearly see the coexistence of the species in this fluctuating environment. Each plot shows one technical replicate of five initial fractions, all of which were drawn from a different biological replicate than in panel **B**. Error bars are the SD of the beta distribution with Bayes' prior probability (see Methods).

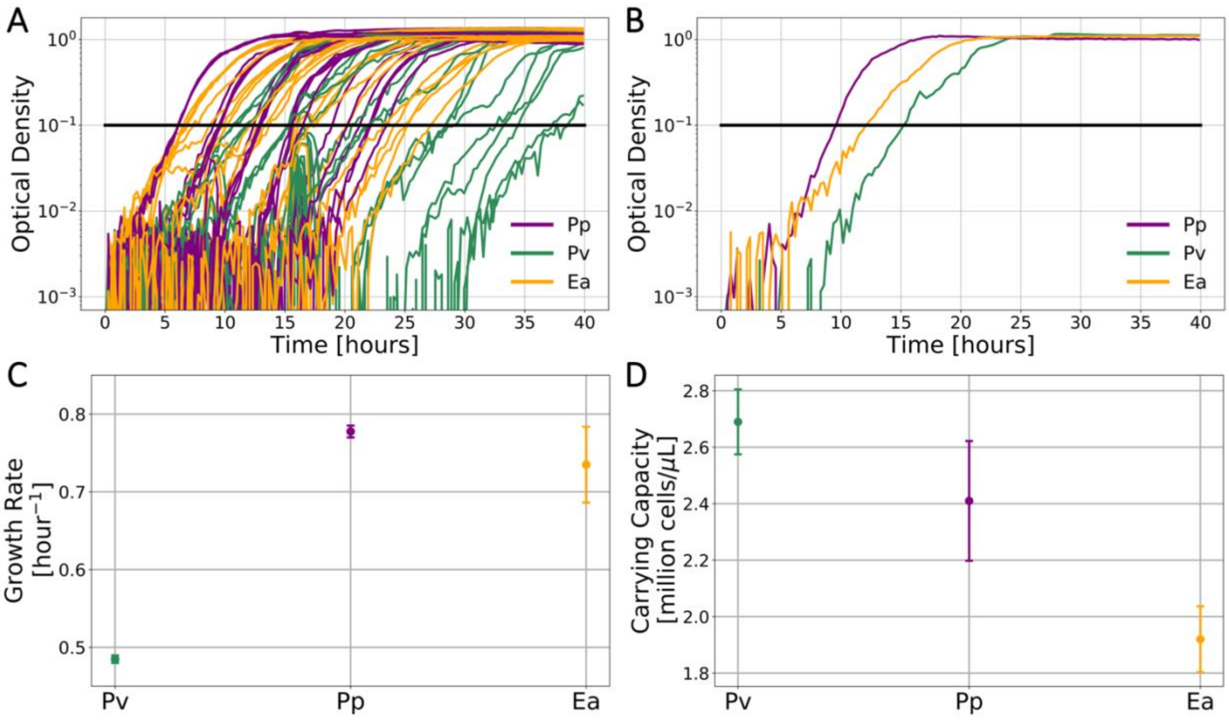


**Supplementary Figure 5: Growth rates for different species were measured using a time-to-threshold method.** To measure growth rates, species were grown in monoculture from a low starting density, with optical density (OD) measured over a period of ~40 hours. Before these measurements, species were grown in 1X LB broth overnight, and then transferred to the experimental medium for 24 hours. The OD of all species was then equalized. The resulting cultures were diluted into fresh medium at factors of $10^{-7}$ to $10^{-3}$. All of the data used to determine the growth rates is shown in **A**, and one set of growth curves is shown in **B**. Background noise has been subtracted from all curves, and no curves have been smoothed. A threshold OD of 0.1 was chosen, and exponential growth was assumed to occur until this threshold. The time each monoculture took to reach this threshold OD was used along with its initial OD to determine the growth rate. By assuming exponential growth to a threshold, we assume no lag time occurs, but the resulting measurement implicitly incorporates lag: longer lag times will cause the measured growth rate to be lower, while shorter lags will have the opposite effect. **C**: Final growth rate measurements were determined for each species by averaging these measurements across all replicates. **D**: Shown are the measured carrying capacities used for simulating the LV model shown in the figures in the main text. Error bars are the SEM of all replicates (n≥16, per species), of which there were four biological replicates and at least four technical replicates of each (plate reader noise due to factors such as condensation caused some replicates to be excluded for some species).

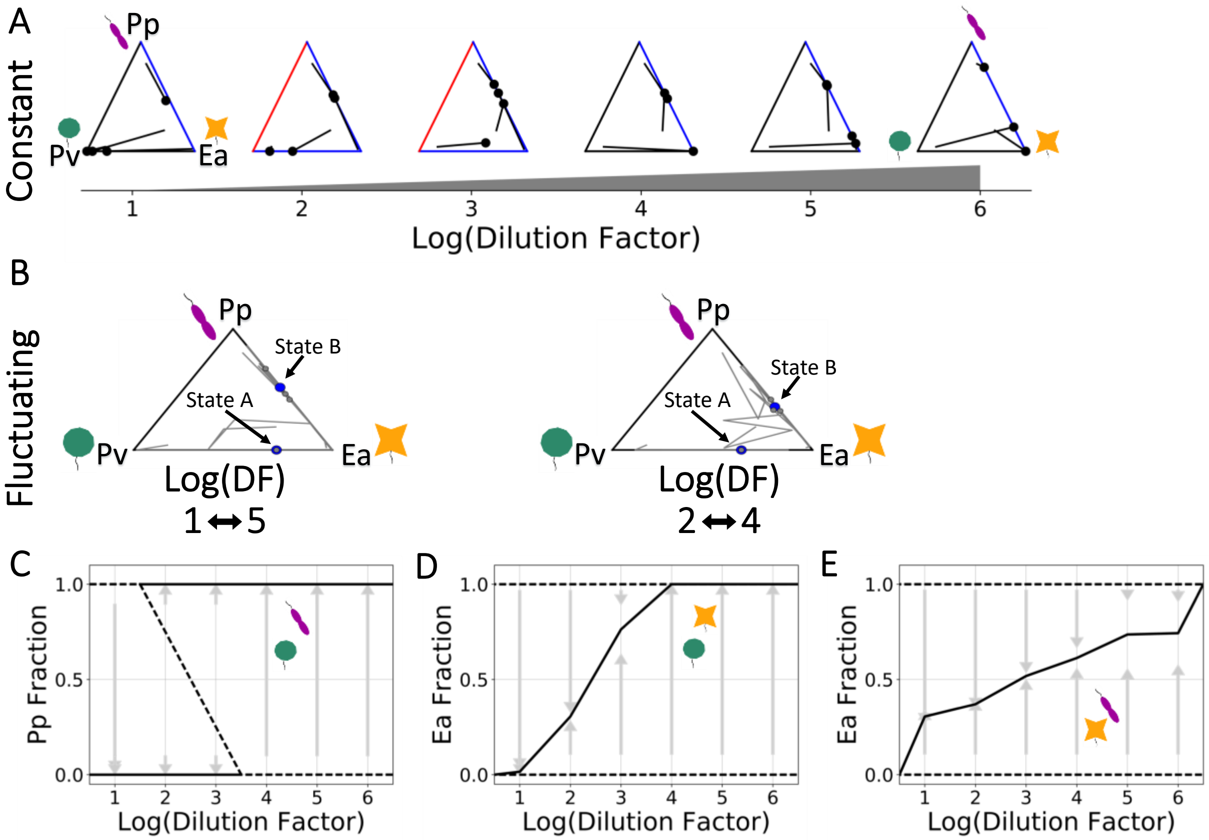


**Supplementary Figure 6: A three-species community reaches similar outcomes in fluctuating and constant environments. A**: Initial and final fractions for each of four starting conditions in each of six constant environments, in which the daily dilution factor (DF) ranged from DF $10^1$ to DF $10^6$. The edges of the ternary plots denote pairwise outcomes; black indicates exclusion, blue coexistence, and red bistability. These outcomes were determined by competing each pair of species in constant environments, the results of which can be seen in the bifurcation diagrams of **C-E**. (Note that one of the pairs, Ea-Pp, violates the model's prediction: although Pp is the faster grower (see Supplementary Figure 5), Ea is slightly favored at higher dilution factors. However, Ea and Pp had the most similar estimated growth rates ($p = 0.17$, compared to $p < 0.01$ for all other pairs of species), which makes the model's prediction more tenuous for this pair.) In **B**, the results of fluctuating the DF in two different regimes are shown, between DF $10^1$ and DF $10^5$, and between DF $10^2$ and DF $10^4$. Both have a constant equivalent DF of $10^3$, and we see that both regimes have the same qualitative outcomes as the constant environment with DF $10^3$, as three initial fractions go to State B and one initial fraction goes to State A, with state labels consistent with Figure 4 in the main text. All plots show data sampled from the same biological replicates.

### Supplementary Discussion

### Supplementary Note 1: Derivation of Lotka-Volterra model modified by added death

The most basic form of the two-species Lotka-Volterra model takes the following form:

$$\frac{\dot{N_i}}{N_i} = r_i - c_{ii}N_i - \sum_j c_{ij}N_j \qquad (3)$$

where $r_i$ is the exponential growth rate of species $i$ (minus any intrinsic death rate), $c_{ii}$ is the rate at which species $i$ inhibits itself, and $c_{ij}$ is the rate at which species $j$ inhibits species $i$. Equation (3) can be re-parameterized to:

$$\frac{\dot{N_i}}{N_i} = r_i\left(1 - \frac{N_i - \sum_j \beta_{ij}N_j}{K_i}\right) \qquad (4)$$

where $K_i = \frac{r_i}{c_{ii}}$ is the carrying capacity and $\beta_{ij} = \frac{c_{ij}}{cii}$ is the competition coefficient. We can further re-parameterize the model by normalizing by carrying capacity:

$$\frac{\dot{\widehat{N_i}}}{\widehat{N_i}} = r_i\left(1 - \widehat{N_i} - \sum_j \alpha_{ij}\,\widehat{N_j}\right) \qquad (5)$$

where $\widehat{N_i} = \frac{N_i}{K_i}$ and $\alpha_{ij} = \beta_{ij}(\frac{K_j}{K_i})$. This version of the model is useful because the competition outcomes depend upon whether the competition coefficients are greater or less than one: stable coexistence occurs when both coefficients are less than one, bistability when both are greater than one, and dominance/exclusion when only one coefficient is greater than one. This leads to the log/log phase space (Fig. 2B, Fig. 3A, Supplementary Figure 7), in which boundaries form where competition coefficients equal one.

The modified Lotka-Volterra model includes an added global death term:

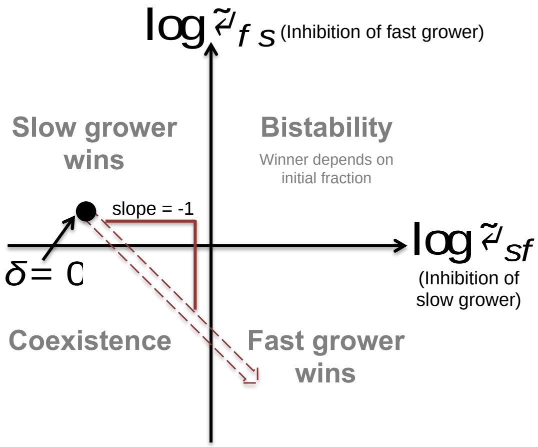


**Supplementary Figure 7:** Re-parameterization of model allows for division of phase space where competition coefficients equal one.

$$\frac{\dot{\widehat{N_i}}}{\widehat{N_i}} = r_i\left(1 - \widehat{N_i} - \sum_j \alpha_{ij}\,\widehat{N_j}\right) - \delta \qquad (6)$$

This term can be absorbed in order to return the model to its previous form (Equation (5)):

$$\frac{\dot{\tilde{N}}_i}{\tilde{N}_i} = \tilde{r}_i\left(1 - \tilde{N}_i - \sum_j \tilde{\alpha}_{ij}\,\tilde{N}_j\right) \tag{7}$$

where $\tilde{r}_i = r_i - \delta,\ \tilde{N}_i = \frac{\widehat{N_i}}{1-\frac{\delta}{r_i}}$ and $\tilde{\alpha}_{ij} = \alpha_{ij}\frac{1-\frac{\delta}{r_j}}{1-\frac{\delta}{r_i}}$. Multiplying $\alpha_{ij}$ by a term means that we add a term to log $\alpha_{ij}$. Due to symmetry, the same term will be subtracted from log $\alpha_{ji}$. As a result, increasing death causes the outcome to move in a line with a slope of negative one through the log/log phase space (Supplementary Figure 7), beginning at the outcome with no added death, (log $\alpha_{ij}$, log $\alpha_{ji}$). If this outcome resides in the quadrant where the slow grower wins, increasing death will eventually result in the fast grower winning. If the trajectory begins in the quadrant where the fast grower wins, however, increasing death will not change the outcome.

While a global death rate of $\delta$ leads to the simple prediction that the fast grower is favored, it is not the most realistic scenario. In reality, different species may be affected by different added death rates. In this case, the expression for the competition coefficients becomes:

$$\tilde{\alpha}_{ij} = \alpha_{ij}\frac{1-\frac{\delta_j}{r_j}}{1-\frac{\delta_i}{r_i}} \tag{8}$$

Taking the log of Equation (8) results in addition of a term to log $\alpha_{ij}$, the same term which will be subtracted from log $\alpha_{ji}$. The outcomes will therefore still move along the same 45° line through the phase space, although not necessarily at the same rate or in the same direction. Added mortality will favor the faster grower if the following condition is met:

$$\frac{\delta_s}{\delta_f} > \frac{r_s}{r_f} \tag{9}$$

We therefore see that the fast grower can still be favored if it is killed at a higher rate (as in the case of $\beta$-lactam antibiotics, which target faster growers by inhibiting cell wall biosynthesis). Furthermore, the growth/competition tradeoff at low dilution is not required to observe outcome changes if the slow grower is selectively targeted; in this case, the trajectory would move from fast grower winning at low mortality, to coexistence or bistability at intermediate mortality, to the slow grower winning at high mortality.

**Supplementary Note 2: Dilution factors can be time-averaged in the Lotka-Volterra model**

As noted in the main text, the Lotka-Volterra model makes an interesting prediction about a fluctuating mortality rate. Because the per-capita growth rates $\frac{\dot{N}_i}{N_i}$ are linear and additive, a fluctuating mortality rate can be time-averaged for the purpose of finding the equilibrium steady state. In fact, even a discrete mortality process, such as the daily dilution factors that we use in experiments, can be time-averaged. The outcome of alternating daily dilution factors (for example, $10^1$ and $10^5$) is the same as the outcome of a constant dilution factor equal to the geometric mean of the two ($10^3$).

To show that the equilibrium state resulting from different dilution factors is equivalent to that from the time-averaged dilution factor, we will begin by examining a single dilution factor. In contrast to a continuous dilution rate, such as in a chemostat, the daily dilution process is itself a fluctuating mortality rate. During most of the cycle, the mortality rate is zero, and the dilution process can be thought of as an instantaneous spike (delta function) in mortality. We will assume that a culture is allowed to grow for a time of length $T$ and then diluted in an instant, whereby the number of cells is divided by the dilution factor $DF$. We can begin by time-averaging the Lotka-Volterra model to model the process:

$$\frac{1}{T}\int_0^T \frac{\dot{N}_i}{N_i} dt = \frac{1}{T}\int_0^T \frac{d}{dt}\log(N_i)\, dt = \frac{1}{T}\int_0^T r_i\left(1 - N_i - \alpha_{ij}N_j\right)dt \tag{10}$$

Additionally, we will assume that the system has reached equilibrium. In this case, $N_i$ grows to the same quantity each cycle, $N_i(T)$, before being diluted to the same quantity, $N_i(0)$. The ratio of the two quantities is equal to the dilution factor:

$$\frac{1}{T}\int_0^T \frac{d}{dt}\log(N_i)\, dt = \frac{1}{T}\log\left(\frac{N_i(T)}{N_i(0)}\right) = \frac{\log(DF)}{T} \tag{11}$$

Now plugging the right-hand side of equation (11) into the left-hand side of equation (10) and defining the time-average $\frac{1}{T}\int_0^T x\, dt = \langle x \rangle$, we can re-write the time-averaged LV model:

$$\frac{\log(DF)}{T} = \frac{1}{T}\int_0^T r_i\left(1 - N_i - \alpha_{ij}N_j\right)dt \tag{12}$$

$$r_i\left(1 - \langle N_i \rangle - \alpha_{ij}\langle N_j \rangle\right) - \frac{\log(DF)}{T} = 0 \tag{13}$$

We removed $r_i$ and $\alpha_{ij}$ from the integrals because they are constants. Equation (13) tells us that at equilibrium (where the per-capita growth rate is equal to zero), the dilution process is equivalent to subtracting a continuous death rate equal to $\frac{\log(DF)}{T}$. A daily dilution will therefore lead to time-averaged population densities that are the same as a continuous dilution rate of this magnitude.

Now that we have shown that, at equilibrium, a daily dilution is equivalent to a continuous dilution rate, we can proceed to show that, at equilibrium, an alternating daily dilution factor is equivalent to a constant time-averaged dilution factor. We will assume that a culture is allowed to grow for a time of length $T$, then diluted by factor $DF_1$, then grown for another time of length $T$, and then diluted by factor $DF_2$:

$$\frac{1}{T}\int_0^T \frac{\dot{N}_i}{N_i}dt = \frac{\log(DF_1)}{T};\ \frac{1}{T}\int_T^{2T} \frac{\dot{N}_i}{N_i}dt = \frac{\log(DF_2)}{T} \tag{14}$$

Time-averaging the entire two-cycle process, we find (dividing each component by 2 to account for the two cycles):

$$\frac{r_i}{2}\left(1 - \langle N_i\rangle_1 - \alpha_{ij}\langle N_j\rangle_1\right) - \frac{\log(DF_1)}{2T} + \frac{r_i}{2}\left(1 - \langle N_i\rangle_2 - \alpha_{ij}\langle N_j\rangle_2\right) - \frac{\log(DF_2)}{2T} = 0 \tag{15}$$

Simplifying Equation (15) reveals that the alternating dilution factor process is equivalent to one with a constant dilution factor equal to the geometric mean of the two:

$$r_i\left(1 - \frac{\langle N_i\rangle_1 + \langle N_i\rangle_2}{2} - \frac{\alpha_{ij}\left(\langle N_j\rangle_1 + \langle N_j\rangle_2\right)}{2}\right) - \frac{1}{T}\log\sqrt{DF_1 DF_2} = 0 \tag{16}$$

To see this equivalence, remember that the constant dilution factor process is modeled with equation (13). Equation (16) can be mapped to equation (13) by redefining the following parameters:

$$DF = \sqrt{DF_1 DF_2};\ \langle N\rangle = \frac{\langle N\rangle_1 + \langle N\rangle_2}{2} \tag{17}$$

A daily dilution regime that fluctuates between dilution factors $DF_1$ and $DF_2$ is thus equivalent, at equilibrium, to one with the same dilution factor everyday equal to the geometric mean of the two factors, $\sqrt{DF_1 DF_2}$.

If the fluctuating environment leads to coexistence of two species, we expect the constant environment to also lead to coexistence of those two species. The same is true of bistability. However, in a finite system, if the system spends too much time in one dilution factor, the outcome may differ from the deterministic solution. For example, suppose species A wins in $DF_1$, species B wins in $DF_2$, and both species coexist in $\sqrt{DF_1 DF_2}$. In a real system, it takes a finite amount of time for species B to go extinct in $DF_1$. If the first cycle is longer than that amount of time, the outcome of the fluctuating environment will be that species A wins rather than coexistence.

The situation becomes even more precarious in the case of bistability: too much time in one dilution factor may move the system to the other side of the separatrix. We thus expect that the location of the separatrix could change when going from a constant dilution factor experiment to an alternating dilution factor experiment, or even that bistability might not result in both regimes. Remarkably, in our experiments, we found that the separatrix was approximately the same in both types of experiments. If we had used an alternation scheme lasting longer than one cycle (for example, two cycles at $DF_1$ followed by two cycles at $DF_2$), it is unlikely that this would still be true.
